# Automated detection of *Loa loa*: a field trial in Cameroon

**DOI:** 10.64898/2026.08.29.745114

**Authors:** Linda Djune-Yemeli, Charles B. Delahunt, Steve Mbickmen Tchana, Jean Gabin Bopda, Yves Aubin Balog, Yannick Yomie Nzeuhang, Dipayan Banik, Matthew D. Keller, Ethan Spencer, María Díaz de León Derby, Zaina L. Moussa, Daniel A. Fletcher, Isaac I. Bogoch, Anne-Laure M. Le-Ny, Joseph Kamgno

## Abstract

Onchocerciasis (river blindness) is targeted for elimination through mass administration (MDA) of ivermectin (IVM) to endemic populations. In areas where loiasis, caused by the blood-borne filarial parasite *Loa loa,* is co-endemic, IVM MDA faced significant challenges because individuals harboring more than 30,000 *L. loa* microfilaria (mf)/mL of blood are at high risk of developing serious adverse events (SAEs) that can sometimes be fatal. An alternative strategy was developed for safe IVM distribution: the so-called “Test and Not Treat” (TaNT), which identifies and excludes individuals with high mf density from IVM MDA and treats only those with minimal risk. TaNT requires a point-of-care diagnostic device to rapidly quantify *L. loa* parasites in field conditions. The NTDscope is a handheld device which captures bright-field videos of whole blood in capillaries. An onboard algorithm detects and counts the live microfilaria of *L. loa* via movement of red blood cells. This paper describes (i) a new detection algorithm; and (ii) a May 2025 field trial in Cameroon using the NTDscope with new algorithm (550 patients). Results for the TaNT use case (i.e. flagging cases with *L. loa* mf densities > 30,000 mf/mL): 94% to 97% sensitivity, and 94% to 97% specificity. The results indicate that the NTDscope and new algorithm provide a greater margin of safety than the predecessor device, and can potentially offer rapid, effective field detection of high mf infections to enable scalability of the TaNT strategy.

## Introduction

Loiasis is an emerging Neglected Tropical Diseases (NTDs) endemic in West and Central African rainforest areas^1,2^. The disease is caused by the filarial nematode *Loa loa*, transmitted between humans through the bite of the tabanid fly *Chrysops*^3^. Epidemiological data estimated that 14.4 million individuals are at risk of infection, with an estimated 10 million cases^2^. Loiasis causes a wide range of clinical symptoms, from no symptoms at all to various severe complications. Notably, infection with *L. loa* is mostly characterized by three mains symptoms: (i) transient angioedema commonly known as Calabar swelling that generally appear on the extremities, and disappear few days later; (ii) ocular migration of the adult parasite under the conjunctiva known as eye worm^3^; (iii) encephalitis, frequent among individuals harboring high microfilariae (mf) density (>30,000 mf/mL)^4^. Furthermore, *L. loa* is responsible for a wide range of atypical clinical manifestations, the more common being recurrent pruritus. Other symptoms involving deep organs (lung, heart, kidney) and excess mortality have been reported among loiasis infected individuals^7,8^. Considering all these, loiasis is under consideration for inclusion in the WHO’s priority list of Neglected Tropical Diseases^19,20,21^.

On the other hand, loiasis has long attracted significant public health attention as an impediment to onchocerciasis elimination. In fact, loiasis is an obstacle to IVM mass administration (MDA) for LF/onchocerciasis in co-endemic areas. Notably, in areas where onchocerciasis and loiasis are co-endemic, individuals harboring more than 30,000 *L. loa* microfilariae (mf)/mL of blood exhibit a high risk of developing serious adverse events (SAEs) including encephalopathy, coma, and death when they are given IVM in the framework of onchocerciasis control^4,10^. To overcome this challenge, an alternative strategy, the so-called Test and Not Treat (TaNT) strategy, was developed. The TaNT strategy uses a mobile-phone-based microscope (originally called the CellScope^11^) to identify and exclude individuals with high *L. loa* microfilarial densities, who are at risk of SEAs, and to treat only those whose mf density falls below the defined safety threshold. This strategy has enabled safe treatment of thousands of individuals during pilot studies conducted in Cameroon.

However, the operational success of the TaNT depends critically on the availability of a reliable point-of-care diagnostic capable of rapidly quantifying *L. loa* microfilarial density in field conditions.

The first device developed for this purpose was the CellScope^11^, consisting of an iPhone-based device plus an algorithm, based on detecting the displacement of blood caused by mf movement. However, continued development necessitated two types of updates: A new device (called NTDscope^18^) was built on an Android platform; and the algorithm was substantially updated to improve robustness.

The goal of this paper is two-fold: (a) Describe the updated algorithm currently on the NTDscope; and (b) Describe this NTDscope’s performance in a recent field trial. The paper is organized as follows: “Methods” first gives a brief overview of the updated algorithm (a detailed walk-through of the algorithm is in the S.I.) and then describes the setup of a May 2025 field trial (hereafter “May25”). “Results” describes algorithm performance results in two settings: (i) on holdout data, assessed in-lab; and (ii) on the May25 field trial, where the updated algorithm was tested as a “black box” diagnostic. This gives two snapshots of the current algorithm’s performance.

## Materials and Methods

This section first briefly outlines some algorithm details, then describes the methods of the May25 field trial.

### 1. Algorithm

#### Goal of algorithm updates

The updates did not alter the basic mechanism of the CellScope algorithm^11^. Rather, they sought to: (i) Adapt the algorithm to Android devices; (ii) Make the algorithm more robust to variability (e.g. replace hard-coded thresholds with adaptive thresholds); (iii) substitute in more effective methods in places; and (iv) tune algorithm hyperparameters using new datasets. The Python codebase can be found at github.com/charlesDelahunt/loaDetectionAlgorithms.

The algorithm does not use Artificial Intelligence. Rather, it uses image-processing and statistical methods to leverage some biophysical priors (e.g. that swimming mf displace red blood cells). Incorporating biophysical priors can (i) improve algorithm robustness by reducing sensitivity to unforeseen variabilities in training and test data, which often impact AI algorithm performance; and (ii) greatly reduce the amount of training data needed, because an AI algorithm does not need to “figure out” the physics of the problem from scratch.

#### Data

The data used to update the algorithm were collected in Cameroon in a 2023 study (hereafter “Cam23”). This data is being released as an open access dataset^22^.

Algorithm development used the sessions from 442 patients of the Cam23 dataset (a session is a set of videos of a single capillary; in Cam23 each patient had 4 sessions). There were two Giemsa reads for each patient, from different blood draws (sometimes discrepant, as expected^14^). The maximum of these two was taken as ground truth Giemsa count.

#### Algorithm brief overview

This subsection gives a few comments of general interest about the algorithm. A detailed walk-through of the algorithm is provided in the S.I.

A capillary filled with fresh, unstained blood provides 7 Fields of View (FoVs), each ≈ 4.5 mm x 6.2 mm. Input to the algorithm is a set of 7 videos, one per FoV. Videos are 5 seconds long. Downsampled frame size is 360 x 480 x 3 with about 13 μm / pixel. *L. loa* mf are themselves invisible in the videos. The algorithm detects them indirectly through the movement of red blood cells as the mf swim (like counting swimmers in a lake via their splashing), by accumulating the motion-induced differences in pixel values frame-to-frame. The algorithm was written and tested in Python^23^, then translated to Android^24^. It estimates a count of active mf/μL in each of the patient’s 7 videos, then from these generates a mf/μL patient count (details in S.I.).

Because the algorithm is known to undercount on high microfilaremia samples (roughly > 15,000 mf/mL), the mf/μL count passes through an adjustment function parametrized to match Giemsa target counts on a calibration set. The final estimated count is given as mf/mL. Patients are flagged as “Not Treat” with ivermectin if the estimated microfilaremia is over 20,000 mf/mL, consistent with field practice. If more than 2 out of 7 videos have insufficient blood (< 80% fill), the test result is rejected.

A word about TaNT thresholds: A standard threshold applied to Giemsa counts to flag “Not Treat” cases is 30,000 mf/mL^4,10^. The CellScope used a threshold of 26,000 mf/mL to add a margin of safety against the risk of device undercount relative to Giemsa. The NTDscope algorithm described here used a threshold of 20,000 mf/mL for the same reason. For both the CellScope and NTDscope, False Positives (FPs) are defined as patients flagged as having device counts above the device threshold but with Giemsa counts below 30,000 mf/mL.

There are two known failure modes:

1. Bulk blood motion in the capillary, perhaps due to capillary leakage, can trigger false positive mf detections. This was seen in some Cam23 videos and is mitigated by a noise floor: Any algorithm count under 750 mf/mL is set to 0 (i.e. putative negative). The theoretical best limit of detection of the NTDscope device, with current capillary design, is about 240 mf/mL^25^. For a detailed discussion of bulk motion and limit of detection, see the S.I.
2. If the capillary is not imaged soon enough, the blood can begin to coagulate, which prevents movement by the mf. This can mask high microfilaremia cases and thus lead to False Negative samples in the TaNT use case. For a detailed discussion of coagulation see the S.I. The protocols for the May25 trial emphasized rapid (< 3 minutes) processing of filled capillaries, which effectively mitigated the coagulation issue.

### 2. May 2025 field trial

#### Ethical considerations

The study received ethical approval from the Centre Regional Ethical Committee for Research on Human Health (CRERSH-Ce) and administrative authorization from the Centre Regional Delegation for Public Health. Communities were informed through the local health system, including district staff, health area personnel, community health workers, and local leaders. In each community, eligible individuals were invited to a central location (health facility or chieftaincy). A team member provided an introductory explanation, which was translated into the local language by community health personnel. All potential participants were given the opportunity to ask questions before deciding whether to participate.

#### Study site and population

The study was conducted in the Okola Health District, located in the Lekie Division of Cameroon’s Centre Region. The district comprises eleven Health Areas. Its low-flowing rivers and forested environment create swampy zones favorable to *Chrysops* breeding. Okola is known to be endemic for loiasis, with an estimated prevalence of about 34% (Djune-Yemeli *et al.,* unpublished). Ivermectin mass drug administration (MDA) began in 1999, but more than 23 cases of encephalopathy were reported during the first round, leading to suspension of MDA across the district. After investigation, MDA was resumed in 5 of the 11 Health Areas, while the remaining six (Okola, Ngoya, Nlong, Mvoua, Lobo, and Ekekam 3) were classified as non-eligible. The present study was carried out in these six non-eligible Health Areas.

#### Enrollment of participants

After the introductory briefing, volunteers proceeded to the registration station where informed consent was obtained. Each participant was then assigned a barcode for identification. They were subsequently directed to the laboratory stations, where they were tested twice using both the NTDscope and the (calibrated thick blood smears) CTBS. Finally, a brief clinical examination was conducted to document any history of Calabar swelling or eye-worm episodes.

#### Samples’ collection and processing

Sample collection focused on performing both the NTDscope test and the CTBS. Notably, rapid diagnosis of loiasis was carried out using the NTDscope on finger-prick blood, following previously described procedures^11^. In parallel, CTBS were prepared using 70 µL of finger-prick blood, following standardized protocols of the ISM laboratory. Briefly, non-heparinized finger-prick blood was collected and drawn onto a microscope slide. The slide was then allowed to dry and stained with 10% Giemsa using standard procedures^12^. On each stained slide, 50 µL (not 70 µL) of blood was examined under a light microscope at 10X objective for blood dwelling mf. All mf present on the smear were counted and reported. Mf of *M. perstans*, also endemic in the study area, were distinguished from those of *L. loa* by size and the absence of a sheath. The final result was expressed in microfilariae per milliliter of blood (mf/mL)^13^.

#### Morbidity assessment

Morbidity assessment focused on detecting the main clinical signs of loiasis, Calabar swelling and subconjunctival migration of the adult worm, using the validated RAPLOA^29, 30^ approach. Briefly, after identification of the local name of eye worm, participants were asked whether they had already experienced it. For those who responded yes, photography was used to confirm their affirmation. Finally, they were asked how long the condition was the last time they had it. Those in which the last eye worm experience did not exceed seven days were recorded as positive for eye worm history. As for the history of eye worm, pictures were used to confirm their affirmations. For Calabar swelling, participants were asked whether they ever experienced swellings under the skin that changed position and disappeared. The medical consultation was conducted in French or English, and when needed, translation was done in local language by a community health worker.

## Results

This section describes the updated algorithm’s performance in two test situations:

1. Validation results for the algorithm using holdout data from the Cam23 dataset^22^. This holdout data was not used to develop or calibrate the updated algorithm, but it came from the same study as the development data.
2. Results from the May25 field trial, in which the updated algorithm was deployed as a black box diagnostic.

### 1. Results on a holdout dataset

This subsection describes algorithm results on holdout data from the Cam23 dataset, calculated in-lab. The result types are: (i) Accuracy relative to Giemsa reads; (ii) False Negative rates in the TaNT context; (iii) Accuracy relative to manual per-video annotations; and (iv) Results of a simulation to assess performance in the TaNT use case.

The count adjustment function (noted in the algorithm overview) was determined on a subset of patients the Cam23 dataset. Results are reported for a holdout set of 1139 sessions from 529 patients (each patient in this holdout has 2 to 3 NTDscope reads due to the data collection protocol).

#### (i) Accuracy vs Giemsa

In general, the current algorithm yields counts that match Giemsa counts with about the same variability of Giemsa counts relative to each other. Patient-level algorithm counts vs max Giemsa for the Cam23 Holdout set are shown in **Fig 1A**. The agreement of the NTDscope to Giemsa is similar to that of one Giemsa read relative to the other (**Fig 1C**). For historical comparison, session-level CellScope counts vs Giemsa are shown in **Fig 1E**. The spread of percentage discrepancies for the NTDscope vs max of two Giemsa reads (**Fig 1B)** is similar to the spread between two Giemsa reads (**Fig 1D**) and the spread of the historical CellScope vs Giemsa (**Fig S4**).

**Fig 1:**
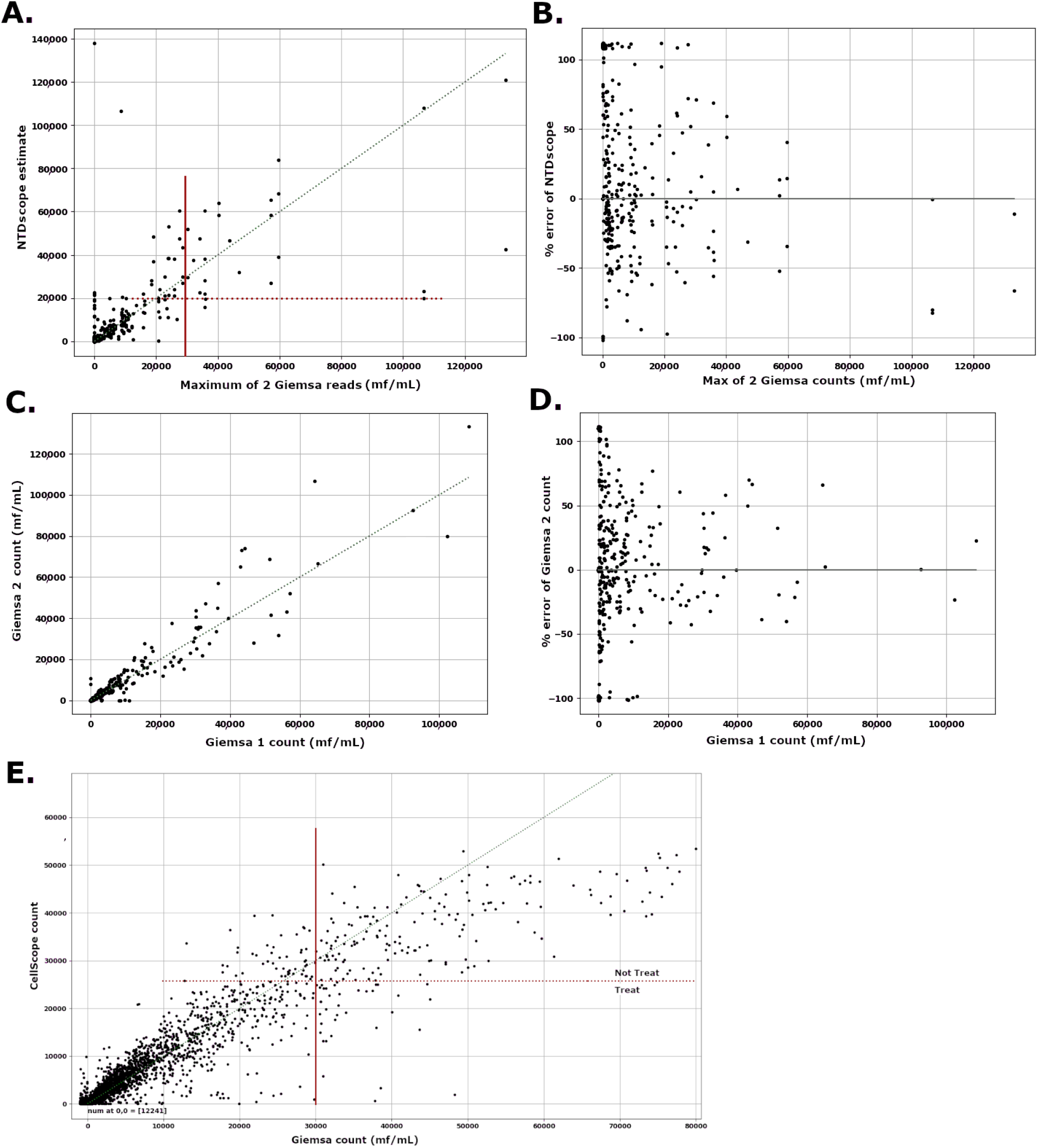
Results on Cam23 Holdout set. **A**: NTDscope count vs Max Giemsa count. There are 631 points at [0,0]. **B**: Percent error of NTDscope vs Max Giemsa count (same data as **A**, visualized differently). **C**: Giemsa 2 count vs Giemsa 1 count. **D**: Percent error of Giemsa 2 count vs Giemsa 1 count (same data as **C**, visualized differently). **E**: Historical data: CellScope vs Giemsa. In **B** and **D**, dot locations are jittered, and errors greater than 100 are bundled at 110%. In **A** and **E**, the lower right quadrant of the red lines contains FNs for the TaNT use case (i.e. undetected high filaremias), while the upper left quadrant contains FPs (i.e. patients erroneously flagged for NT). The horizontal red lines are device thresholds (20,000 for NTDscope, 26,000 for CellScope). **Alt Text:** Five scatterplots which show that Giemsa reads, CellScope, and NTDscope all have similar spreads of error relative to a Giemsa ground-truth.

#### (ii) False Negatives in the TaNT context

A crucial measure of effectiveness in the TaNT use case is the rate of False Negatives (FNs). An FN in the TaNT context is a patient with microfilaremia > 30,000 erroneously labeled as low microfilaremia and therefore eligible for ivermectin, potentially creating risk of an adverse event. FN rate can be calculated empirically as **Alt Text:** Equation for FN rate, which is the percentage of patients with over 30,000 mf/mL Giemsa counts for which the device count is below the device cutoff threshold.

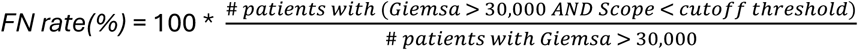

The FN rates for CellScope and the updated NTDscope are shown in **Table 1**. The FN rate for the NTDscope as calculated on the Cam23 Holdout varies according to whether one, or the other, or the maximum of both Giemsa reads are used as the reference. This highlights the impact of the choice of reference. NTDscope FN rates from the May25 trial are discussed below, in “Results for the May25 field trial”.

**Table 1:** FN rates in the TaNT context. “# patient reads” means the number of patients with Giemsa counts > 30,000.

| Device (dataset) | Reference | # patient reads | FN rate (%) |
| --- | --- | --- | --- |
| CellScope (from ref<sup>3</sup>) | Single Giemsa | 245 | 15.9 |
| NTDscope (Cam23) | Max of 2 Giemsas | 26 | 7.7 |
| NTDscope (Cam23) | 1<sup>st</sup> Giemsa only | 26 | 7.7 * |
| NTDscope (Cam23) | 2<sup>nd</sup> Giemsa only | 26 | 11.8 * |
| NTDscope (May25) | Single Giemsa | 33 | 2.9 to 6.1 |

\* The FN rates vary because the two Giemsa reads often differ. For a given patient, either 1<sup>st</sup> or 2<sup>nd</sup> Giemsa can be higher (as seen in **Fig 1D**).

The FP rate (i.e. cases with < 30,000 mf/mL but labeled as unsafe to treat) was 2.3% for the updated algorithm, vs 0.3% for the CellScope. Part of this increase may be due to a higher fraction of Giemsa-positive cases in Cam23 (31% of Cam23 sessions were positive, vs 18% of CellScope’s).

#### (iii) Accuracy vs manual video annotation

Humans can review videos to annotate mf movement, providing a video-specific ground truth distinct from Giemsa counts (Giemsa counts are patient-level, not video-level, and are made on separate blood samples). The algorithm’s raw mf counts per video generally match well to human counts per video (**Fig S2**), indicating that the algorithm is keying off of microfilaria movement. Some points to note: (i) The algorithm has many false positive counts (due to bulk motion which humans readily identify); (ii) the spread above 35 mf/video is likely due to human error as well as algorithm-human disagreement, since manual counting above 35 mf is difficult.

#### (iv) Simulations to estimate FN rate

The accuracy of the algorithm’s FN rates in Table 1 is limited by the relatively low number of high-count cases in our datasets. Therefore, we ran a simulation, separately for NTDscope on Cam23 Holdout and for CellScope data^3^, as follows:

1. Define 20,000 samples with Giemsa counts uniformly distributed from 30,000 to 50,000 mf/mL.
2. For each sample, simulate the algorithm’s estimated count as follows: (i) characterize the statistical distribution of the algorithm’s percentage errors on the actual higher-microfilaremia cases (> 20,000 mf/mL) in the dataset; (ii) for each sample, randomly draw from this distribution to assign a percentage error to modify the sample’s Giemsa count. This gives a simulated set of Giemsa readings with algorithm estimates, whose errors have the same statistical properties as the empirical errors.
3. Calculate FN rates for windows with width 2,000 mf and with centers sweeping through the true Giemsa values (31,000 to 49,000), using the algorithm’s “NT” threshold (20,000 for NTDscope and 26,000 for CellScope) to determine FN status This gives a curve of estimated FN rates for various Giemsa counts.

The simulated FN rates, for the NTDscope with updated algorithms and for the CellScope are shown in **Fig 2**. NTDscope’s FN rates vary from 1.5% (for microfilaremias near 50,000) to 7.5% (for filaremias near 30,000). By contrast, CellScope’s FN rates range from 5% to 18%. This suggests that the NTDscope with updated algorithms improves the margin of safety.

**Fig 2:**
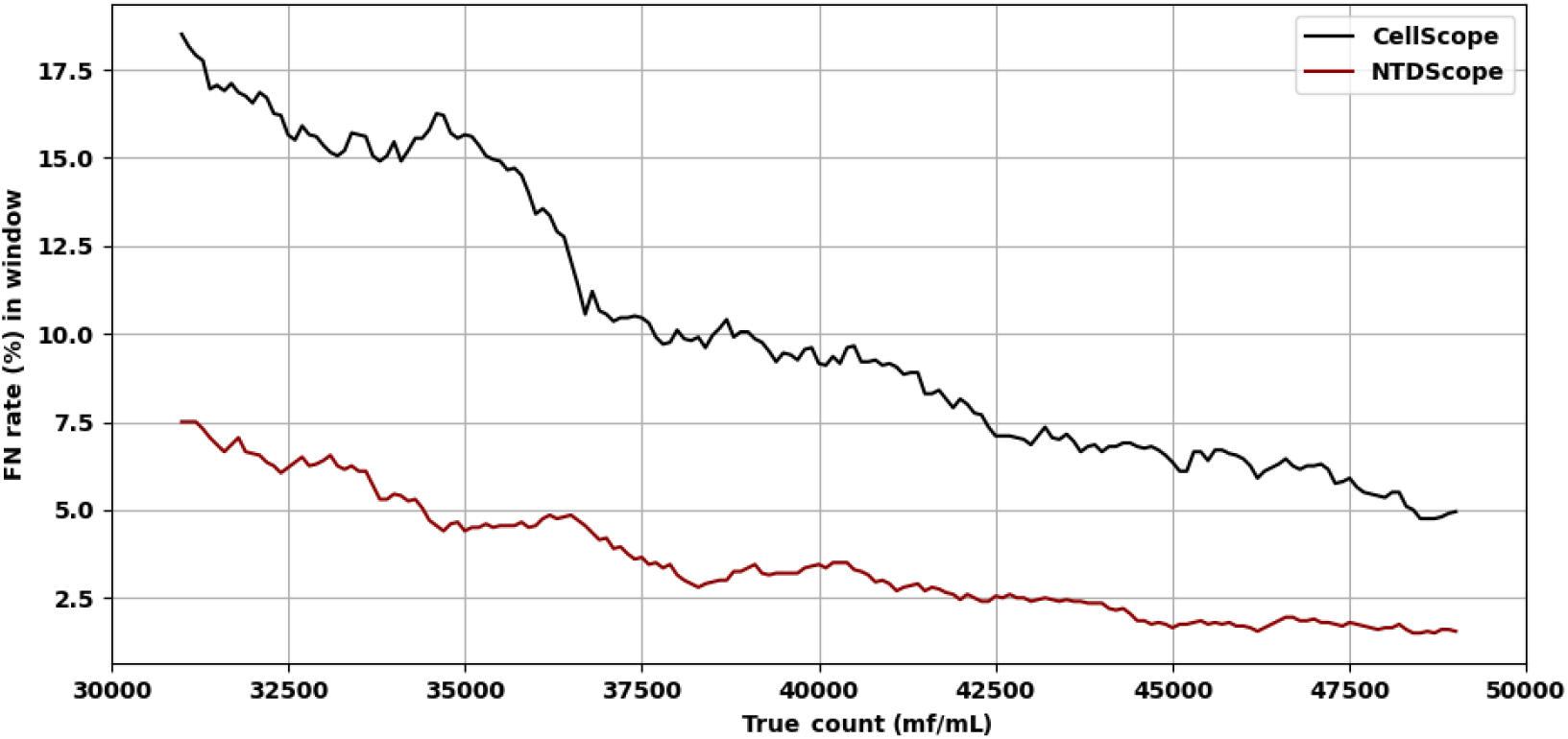
Simulation results for CellScope (black, top) and NTDscope (red, bottom): FN rates for a rolling window, 2000 mf/mL wide, vs the Giemsa count at window center. For example, the FN rates for cases with counts between 39,000 and 41,000 are ≈ 3.5% for NTDscope and ≈ 9% for CellScope. **Alt Text:** Plot of a simulation, the NTDscope has much lower FN rates than the CellScope.

### 2. Results for the May 2025 field trial

The updated algorithm, loaded onto the NTDscope, was deployed in a field trial in Cameroon in May 2025, giving results for 550 patients. Each patient had one Giemsa read and two reads by separate NTDscopes (on different capillaries). There were two processing sites, each with their own pair of NTDscopes and distinct patients. For ease of reporting, the results of one of the NTDscopes from Site 1 are concatenated with results of one of the NTDscopes from Site 2 (arbitrary choice), and similarly with the remaining two NTDscopes. These combined NTDscope results are reported as NTDscope “A” and NTDscope “B”. A and B each have one result per patient. **Table 2** gives statistical results for three use cases: (i) TaNT, for which a “positive” is a patient with Giemsa count > 30,000 mf/mL; (ii) a standard positive vs negative, for which a “positive” is a patient with Giemsa count > 0 mf/mL; and (iii) “greater than 1200 mf/mL”, for which a “positive” is a patient with Giemsa count > 1200 mf/mL (the purpose of this last use case is to highlight the effective limit of detection; results showed that the algorithm was very accurate distinguishing cases as above or below 1200 mf/mL, but had poor specificity detecting cases with 0 mf/mL).

**Table 2:** Statistical results for the two NTDscopes, for three use cases. All results are nearest integer percentages (%).

| Statistic | NTDscope A | NTDscope B |
| --- | --- | --- |
| **TaNT (> 30,000 mf/mL)** |  |  |
| Sensitivity (% correctly flagged as “NT”: TPs) | 94 | 97 |
| Specificity (% correctly flagged as “T”: TNs) | 97 | 94 |
| **Pos/Neg (> 0 mf/mL):** |  |  |
| Sensitivity | 92 | 91 |
| Specificity | 66 | 71 |
| **>1200 mf/mL:** |  |  |
| Sensitivity | 98 | 99 |
| Specificity | 94 | 94 |

Overall, the algorithms had high accuracy vs Giemsa counts > 1200 mf/mL, similar to its performance on the Holdout (as described above). Scatterplots of NTDscope counts vs Giemsa are given in **Fig 3A** (for A) and **Fig 3B** (for B).

**Fig 3:**
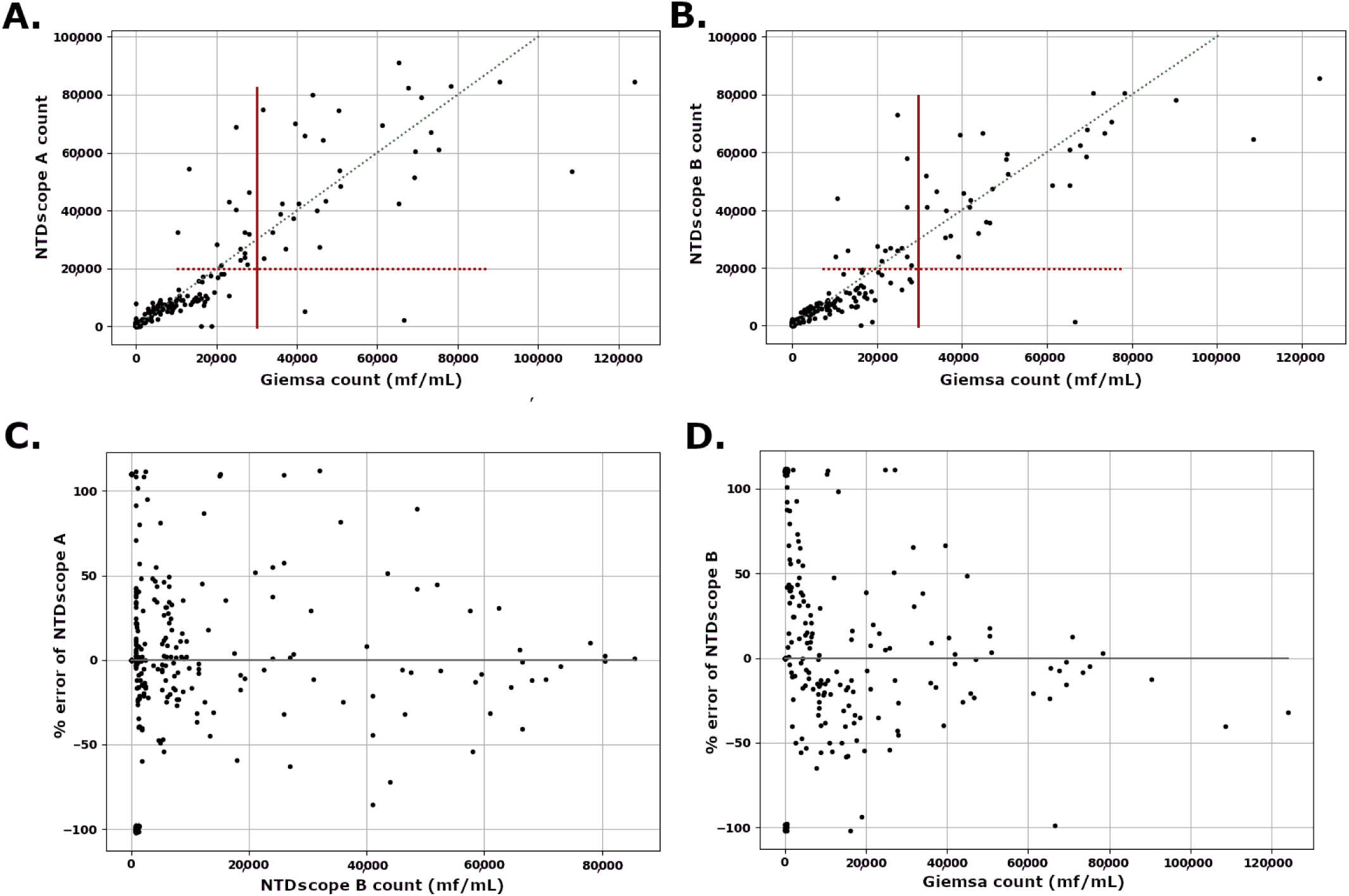
Results on the May25 trial. **A**: NTDscope “A” vs Giemsa counts for 550 cases. There are 233 cases at [0,0], and two False Negatives (in the TaNT sense, at red arrows). **B**: NTDscope “B” (y-axis) vs Giemsa (x-axis) counts for 550 cases. There are 250 cases at [0,0] and one False Negative (at red arrow). %. **C**: Inter-device comparison: Percentage discrepancy of NTDscope A (y-axis) vs NTDscope B as reference (x-axis). The inter-device discrepancy is similar to that of 2 Giemsa reads in Fig 1D. **D**: Percentage error of NTDscope B (y-axis) vs Giemsa as reference (x-axis). This subplot revisualizes the data of subplot B. The error plot of NTDscope A vs Giemsa is very similar (not shown). Dot locations are jittered, and errors greater than 100% (including positive counts on negative samples) are bundled at 110. **Alt Text:** Four scatterplots showing that the counts of both NTDscopes are well-correlated with Giemsa counts, that FNs are very rare, and that interdevice agreement matches device-Giemsa agreement and interGiemsa agreement.

The spread of percent error of NTDscope vs Giemsa (**Fig 3D**) and the inter-NTDscope discrepancies (A vs B, **Fig 3C**) were similar to both the measured inter-Giemsa reader discrepancy for Cam23data (**Fig 1D**) and the percent error of NTDscope vs Giemsa for the Cam23 Holdout (refer **Fig 1B**). The FN rate for TaNT (i.e. percentage of patients with Giemsa count > 30,000 but flagged as eligible for ivermectin by NTDscope) was 6% (NTDscope A) or 3% (NTDscope B), lower than both CellScope and the NTDscope on the Cam23 Holdout. The FP rates were higher in the May25 trial than on the Cam23 Holdout (2.9% and 6.2%, vs 2.3%), while the proportion of Giemsa-positive patients was similar (36% vs 31%).

## Discussion

This study demonstrates that the updated NTDscope algorithm represents a substantive advance over the original CellScope platform for point-of-care quantification of *Loa loa* microfilaremia, particularly in the context of the Test-and-Not-Treat (TaNT) strategy. The transition from an iPhone-based system to an Android platform substantially improves scalability and cost-effectiveness, both of which are critical considerations for large-scale deployment in loiasis–onchocerciasis co-endemic regions. In parallel, algorithmic updates designed to improve robustness under field conditions have translated into improved diagnostic performance in real-world use.

From a programmatic perspective, the most important finding is the reduction in False Negative (FN) rates in the TaNT safety-critical range (>30,000 mf/mL). Because FN errors in this context correspond directly to individuals at risk of ivermectin-associated serious adverse events, the observed decrease in FN rates relative to the CellScope represents a meaningful gain in safety margin. This improvement was evident both in retrospective analyses using holdout data, and during the May 2025 field trial in Cameroon, where the algorithm was deployed as a black-box diagnostic under operational conditions. While the updated algorithm exhibited modestly higher False Positive (FP) rates resulting in a small proportion of individuals being unnecessarily excluded from ivermectin treatment, this trade-off is generally acceptable in the TaNT framework, where prioritizing safety over maximal treatment coverage is a design principle.

The agreement between NTDscope estimates and calibrated thick blood smear (Giemsa) counts was comparable to the inter-reader variability observed between independent Giemsa readings. Similarly, inter-device variability between separate NTDscope units was of the same order as discrepancies between Giemsa reads performed on different blood samples. These findings underscore that much of the observed variability reflects intrinsic biological heterogeneity and reference-standard imprecision rather than device-specific limitations. In this sense, the NTDscope performs at a level consistent with established parasitological methods, while offering substantial advantages in speed, portability, and field suitability.

Beyond its immediate application to TaNT, the current NTDscope algorithm provides a strong foundation for broader use cases, including *Loa loa* prevalence mapping. Rapid, quantitative, point-of-care measurements could support more granular spatial risk stratification and improve modeling efforts aimed at delineating areas where ivermectin MDA can be safely implemented. Such capabilities are particularly relevant in historically excluded districts, where uncertainty regarding loiasis risk has stalled onchocerciasis elimination efforts.

The fact that the algorithm’s performance on the May25 trial data matched or even exceeded its performance on the Cam23 Holdout set is somewhat noteworthy, since algorithm performance typically deteriorates as the test sets move further away from the training set (“distribution shift”^31^). We see two reasons for the consistent high performance. First, in the light of the coagulation issues uncovered in the Cam23 trial (due to running some capillaries twice), the protocol for the May25 trial was amended to emphasize rapid processing of capillaries. This resulted in markedly less coagulation. Second, the algorithm may simply be robust, perhaps because it is based on physical properties of the use case. Complex models with high free parameter counts are sometimes more brittle than simple models^9^.

Despite these advances, several limitations and opportunities for improvement warrant consideration. First, undetected blood coagulation remains a potential source of FN results, as coagulated samples can suppress microfilarial movement and thereby evade motion-based detection. Although adherence to strict field protocols (e.g. processing capillaries within three minutes of blood collection) substantially mitigated this risk during the May 2025 trial, the incorporation of an automated coagulation-detection module would provide an additional layer of safety. Such a feature could identify compromised samples in real time and prompt retesting, further reducing the likelihood of clinically consequential FN errors.

Second, results from the field trial suggest that the effective lower limit of detection (LoD) of the current system lies between approximately 750 and 1,200 mf/mL (seen in Table 2, in the improved specificity when using a 1200 mf/mL threshold). Improving sensitivity at low microfilaremia levels could enhance field acceptance by reducing discordance between clinician expectations and device output, particularly in cases of visibly low-level infection. Greater precision in this range may also improve the performance of prevalence-mapping models. In addition, a lower LoD would expand the utility of the NTDscope to other filarial diseases, such as lymphatic filariasis or *Mansonella* species, which often present with lower peripheral microfilarial densities. The present LoD is largely constrained by identifiable failure modes, most notably bulk blood motion, likely due to capillary leakage, and flickering illumination, which generate spurious motion signals. Importantly, these failure modes are well characterized, and algorithmic safeguards against them appear feasible; preliminary work suggests that machine-learning–based classifiers could effectively identify and suppress such artifacts.

Third, while the present analysis focused on the widely used 30,000 mf/mL threshold relevant to severe adverse event risk, emerging evidence suggests that lower thresholds (e.g. around 8,000 mf/mL) may be clinically relevant in certain subpopulations. Evaluation of NTDscope performance relative to alternative safety thresholds was beyond the scope of this study but represents an important direction for future work, particularly as TaNT strategies continue to evolve.

Lastly, the current algorithm does not distinguish among co-endemic filarial species. In regions where *Loa loa* and *Mansonella perstans* overlap, species-level differentiation would enable more precise epidemiological studies and broaden the NTDscope’s relevance to other neglected tropical diseases. Enhancing species discrimination would further strengthen the device’s value as a multipurpose field diagnostic platform.

The updated NTDscope algorithm demonstrates improved performance and potential scalability relative to its predecessor. These advances support its continued development and deployment for TaNT-based ivermectin distribution and for *Loa loa* surveillance more broadly. With further refinements addressing coagulation detection, low-density sensitivity, and species differentiation, the NTDscope has the potential to play a significant role in overcoming longstanding diagnostic barriers to filarial disease control and elimination in Central and West Africa.

## Acknowledgements

This work was funded by Global Health Labs, Inc.

## Conflicts of interest disclosure

IIB consults to the Weapons Threat Reduction Program at Global Affairs Canada.

## Supplementary Information

### Algorithm details

This appendix provides a more detailed walk through the algorithm. It also discusses limit of detection and known failure modes (bulk motion, coagulation).

#### Algorithm input

A capillary filled with fresh, unstained blood provides 7 non-overlapping Fields of View (FoVs). Input to the algorithm is a set of 7 videos, one for each FoV. Videos are 5 seconds long, captured at 15 to 35 frames per second (FPS). FoV size is ≈ 4.5 mm x 6.2 mm. Downsampled (by 3x) frame size = 360 x 480 x 3 with pixel pitch ≈ 13 μm / pixel.

Technical comments: (i) A slight overlap of or missed space between FoVs does not change the result. (ii) We don’t know how FPS < 15 would perform. (iii) Downsampling frame size by 3x greatly sped up processing and did not reduce accuracy, while downsampling by more than 3x degraded accuracy. (iv) Each video gets divided into five 1-second segments: having several segments is valuable for noise reduction, though we don’t know the constraints on segment duration and count parameters.

#### Algorithm steps

*For each video in a patient’s capillary:*

1. Extract the stack of downsampled green channel images (360 x 480 x 1). The green channel functions as a greyscale. Also extract a single frame’s red channel image.
2. Identify a mask of the blood-filled region, using the ratios of pixel intensities of a frame’s red and green images. This is the algorithm’s only use of color.
3. Comment: The NTDScope’s centrally located illumination causes a dome-shaped variation in pixel intensities. However, luminance normalization is expensive and did not improve algorithm performance because the algorithm tallies differences between frames, not absolute intensity values.
4. Divide the greyscale image stack into 5 non-overlapping 1-second segments, using the FPS provided in the video metadata.
5. *For each 1-second segment in the video*:

a. Calculate a sequence of "average images" *F(i)* that each average 5 consecutive frames. Each *F(i)* steps forward by 1 frame. This step mitigates high-frequency noise. The “5” parameter implicitly assumes FPS >= 15.
b. Create a "difference image" *D*, where each pixel of *D* is the cumulative sum of absolute differences between consecutive *F*’s. So *D* is bright where the pixels values change a lot between consecutive *F(i),* and *D* is dark if the pixels do not change (e.g. in stationary blood). Large pixel differences are caused by:

i. Swimming mf: Displacement of red blood cells gives localized pixel differences along their movement paths.
ii. Distractor phenomena: (1) Bulk motion of blood due to a leaky capillary gives large-region pixel differences; (2) flickering illumination can give large-region pixel differences; and (3) camera jitter can give pixel differences at capillary borders. Later steps seek to identify these false alarms.
c. Normalize *D* by the number of frames. Also do some blurring and eroding steps to remove small noise regions while retaining and separating larger bright regions (assumed to be mf splashes).
d. Create a "z-score image" *Z*, where each pixel is the z-score of its value in *D* (Mahalanobis distance, i.e. the number of standard deviations away from the median value of *D* it is). For this we use the right sided std dev from an asymmetrical std dev calculation^25^. We now have two images, *D* and *Z*, for each 1-second segment (**Fig S1**).

**Fig S1:**
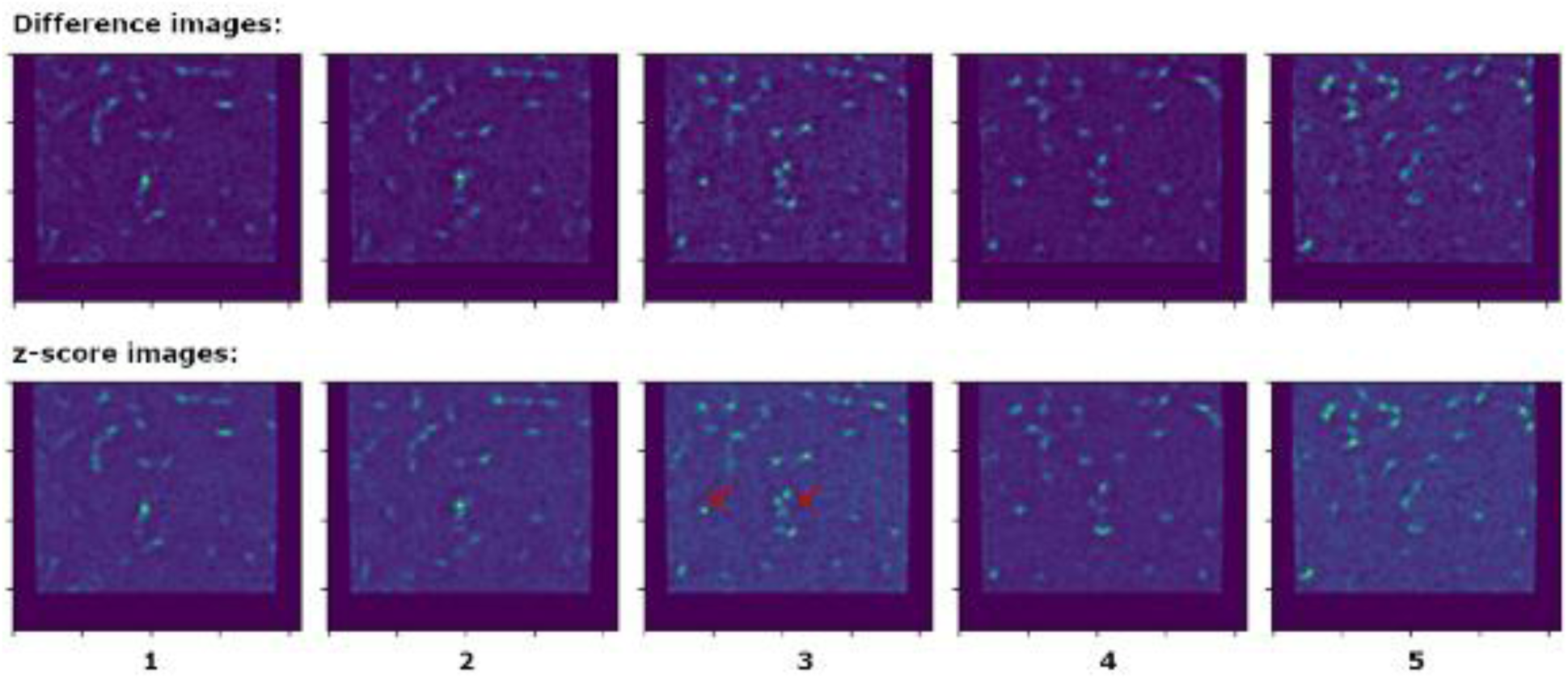
Difference (top) and z-score (bottom) images (D and Z) for five 1-second segments. Each column is for a 1-second segment. Red arrows indicate mf paths visible in the central region of segment 3 that are not visible in other segments. **Alt Text:** Heat maps for each 1-second segment, showing mf locations as bright regions.
6. Search for microfilariae: For each 1-second segment: First mask both *D* and *Z* with the blood mask and slightly erode the mask to avoid effects due to the capillary boundary, to give a region of blood pixels to examine. Then apply thresholds to D and Z to get binary images of likely splashes. Then apply blob detection to the *AND* of the two binary images. Label the centroid of each sufficiently large blob as a mf location. Comments: (i) Each mf is labeled by a single *x-y* coordinate pair, even if it has a long travel path (some mf are more athletic than others); (ii) The *AND* condition on the two binary images says that to be a “mf pixel”, a pixel must have sufficiently bright raw value (*D*) *and* must also be extraordinarily bright relative to other pixels (*Z*). Ideally, the *Z* condition filters errors due to high or low raw illumination, while the *D* condition filters errors due to noisy background signal.
7. Given mf locations in each 1-second segment, get a mf count for the full video: Sometimes mf only appear in a subset of the one-second segments (noted by red arrows in **Fig A1**). Therefore we take the union of mf locations in the various segments, by counting up all mf locations from all segments that are sufficiently far from any other mf location.
8. Repeat search (steps 6 and 7 above) if needed: The first Microfilaria Search uses a pair of high (strict) thresholds. But the algorithm is known to undercount on high mf segments. So if the resulting Union Count is above a fixed value, then the microfilaria search and union count are rerun using slightly lower thresholds, to catch more mf. If this resulting union count is above another fixed value, then the microfilaria search and union count are rerun again, with a lower pair of thresholds. The mf locations from the final search are returned.
9. Scale the mf count according to estimated volume of blood searched to calculate the video’s mf count per µL.

Given the per µL estimates per video, the patient’s mf count per µL is set equal to the second highest of their 7 video counts. This simple method aims towards a high estimate while reducing noise from videos with outlier counts. The reported mf/mL is then a piecewise linear adjustment of the per µL estimate, to account for the known fact that the algorithm (and human annotators) undercount mf relative to Giemsa counts at higher filaremias.

The algorithm’s adjustment function deployed in the May25 trial was different than the one applied to the Cam23 Holdout. The Cam23 Holdout adjustment function, when applied retroactively to the May25 data, yielded almost identical key statistics (FN and FP rates, etc), but overcounted at high filaremias (> 20,000 mf/mL).

#### Consistency with manual video counts

As noted in the main text, the algorithm’s counts per video closely match human counts, as seen in

**Fig S2**. Note, however, the many false positive counts due to bulk motion, seen in the lower left.

#### Limit of Detection

The theoretical best limit of detection (LoD), assuming perfect mf detection with no errors, is governed by Poisson statistics of rare events that determine the probability of at least one live mf existing in the capillary. This in turn depends on the volume of blood sampled, with smaller samples giving higher LoDs^25^. The NTDScope images about 25 µL of blood (7 videos, each imaging about 3.75 µL of blood). Poisson distribution calculations^25^ yield that to ensure at least one mf in 25 µL of blood, filaremia inside the capillary must be at least 120 mf/mL. However, mf counts inside the capillary are empirically only about 50% of counts outside, perhaps due to a sieving effect at the capillary entrance. So the theoretical best LoD is about 240 mf/mL.

**Fig S2:**
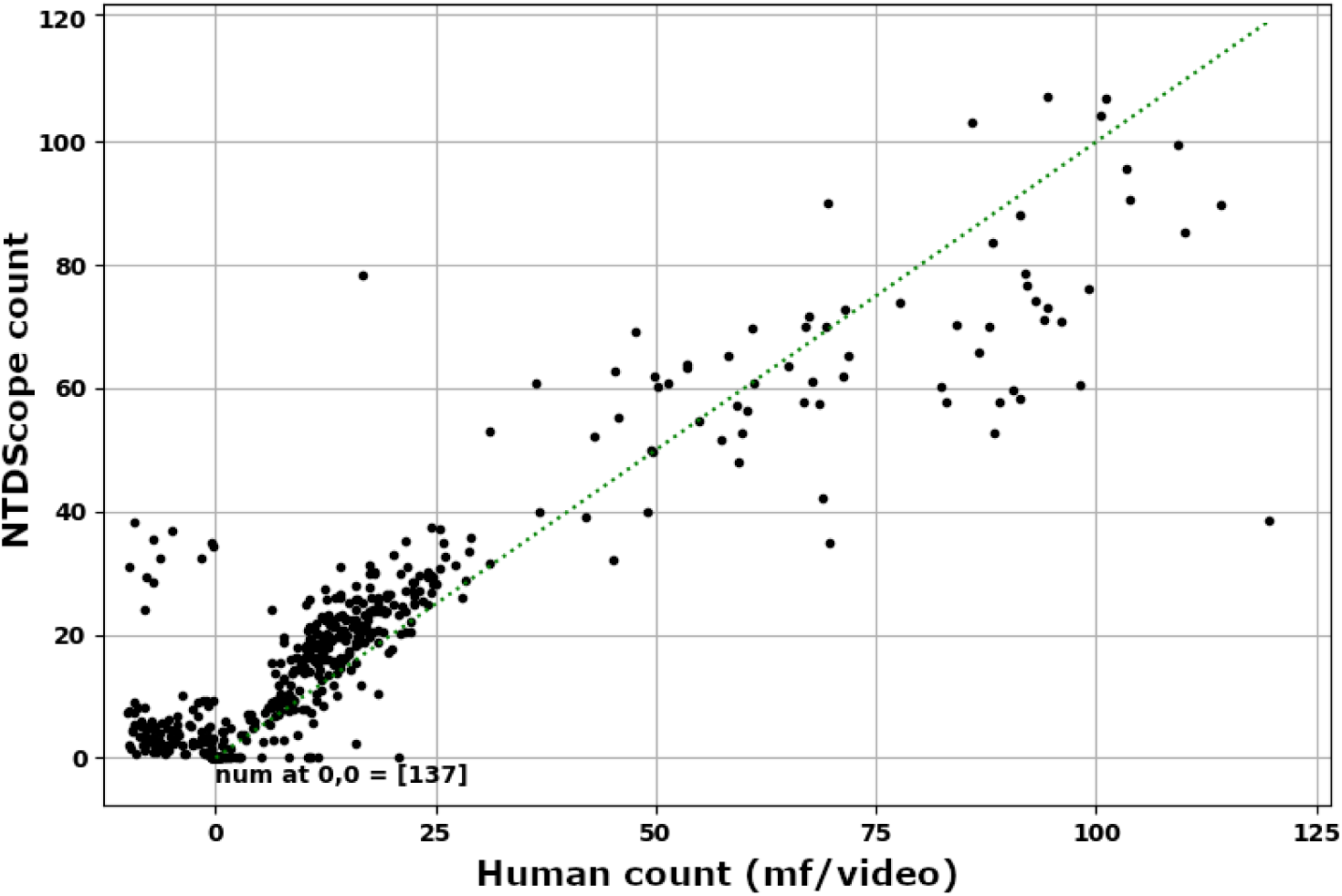
Algorithm mf counts (y-axis) vs human mf counts (x-axis) per video. Cam23 data. Dots for videos with human counts = 0 are spread out between 0 and -15 to better show the algorithm’s false positive counts. **Alt Text:** A scatterplot showing that algorithm counts match human counts per video.

#### Known failure modes

Certain issues can either (1) trigger erroneous mf detections, or (2) mask high filaremia cases:

1. **Mistaken mf detections**: Ideally, the blood in the capillary is motionless except when disturbed by swimming mf, so the background of the difference images will be nearly zero. However, in the Cam23 dataset several videos contained regions of flowing blood (“bulk flow”) perhaps due to leaky capillaries. This flow increases the movement captured in the difference images and causes erroneous mf locations. In a similar effect, bumping the device during imaging can induce bulk motion (a “ripple-in-pond” effect). In a few cases, videos appeared to have a flickering effect, presumably due to illumination flicker. This can cause pixel value differences and thus apparent motion to be captured in the difference images, giving erroneous mf locations.
2. **Coagulation** can lead to False Negative samples in the TCNT use case: If the capillary is not imaged soon enough, the blood can begin to coagulate, which prevents movement by the mf. A common visual difference between coagulated and liquid samples is, roughly speaking, that liquid samples have a more uniform orange color, while coagulated samples have a more “checkerboard” pattern of white and red, since as the blood coagulates it pulls away from small regions in the capillary (see **Fig S3**). We developed a coagulation classifier as described below had holdout sensitivity ∼60% and specificity ∼90%. Since this accuracy was not sufficient for deployment, protocols for the May25 trial emphasized rapid (< 3 minutes) processing of filled capillaries. The importance of mitigating this issue, either by an algorithm module or protocols, is addressed in the Discussion.

The Coagulation Classifier had as input the green channel (and optionally the red channel also) of a single RGB frame per video. The features were Fourier spectra (to capture the checkerboard pattern), and optionally pixel intensity histogram bin counts (to distinguish dark-light vs uniform orange patterns). These were fed into a Support Vector Machine classifier. Ground truth were manual per-video annotations on a train-val set of 62 complete sessions (434 videos) as described in [22], divided into 5-fold train-val splits stratified by session. For more details see [32]. Experiments with DNNs had inferior performance to the above classifier. Because accuracy on the 5-fold validation set was insufficient, no holdout set was assembled.

**Fig S3:**
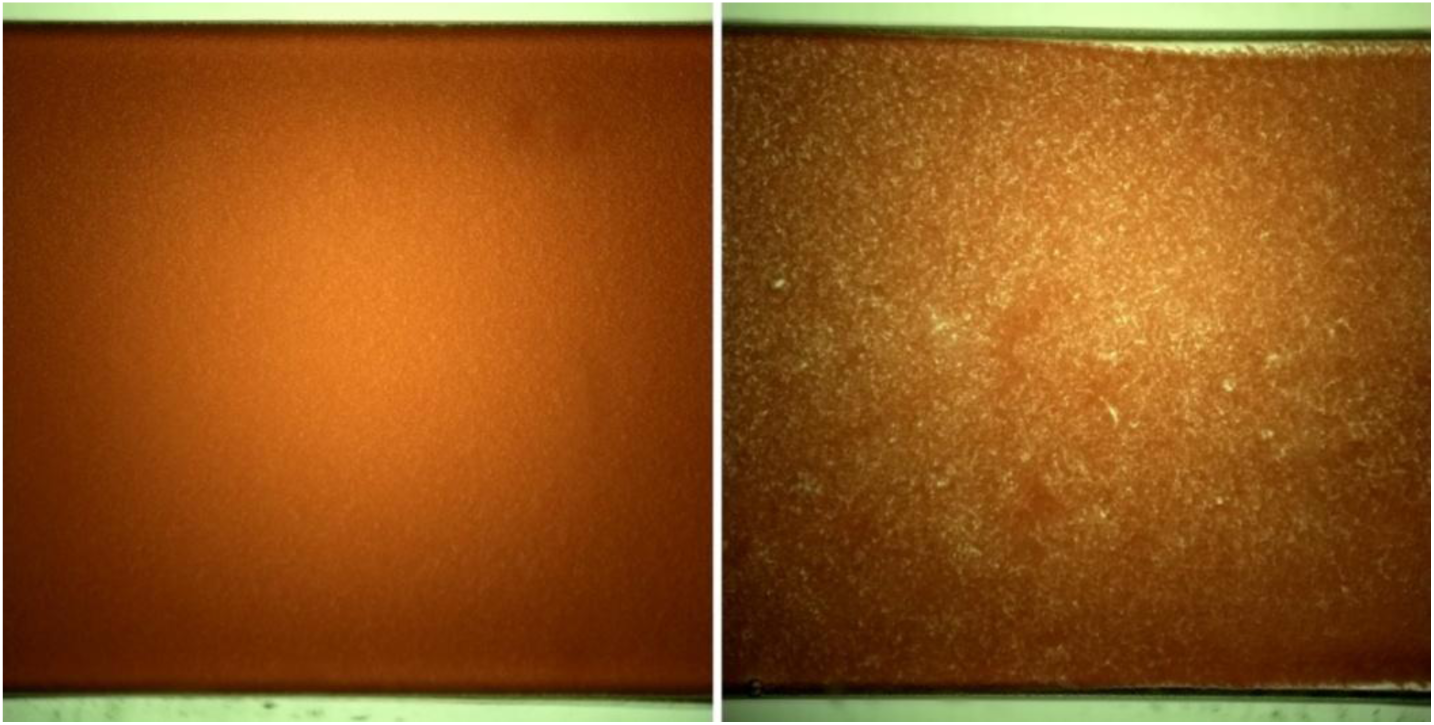
Normal (left) and heavily coagulated (right) blood samples. **Alt Text:** Paired video frames showing uncoagulated vs coagulated blood.

#### CellScope error relative to Giemsa counts

As noted in the main text, the percentage error of NTDscope relative to Giemsa counts (main text, **Fig 1B**) is very similar to that of the historical CellScope relative to Giemsa (**Fig S4**).

**Fig S4:**
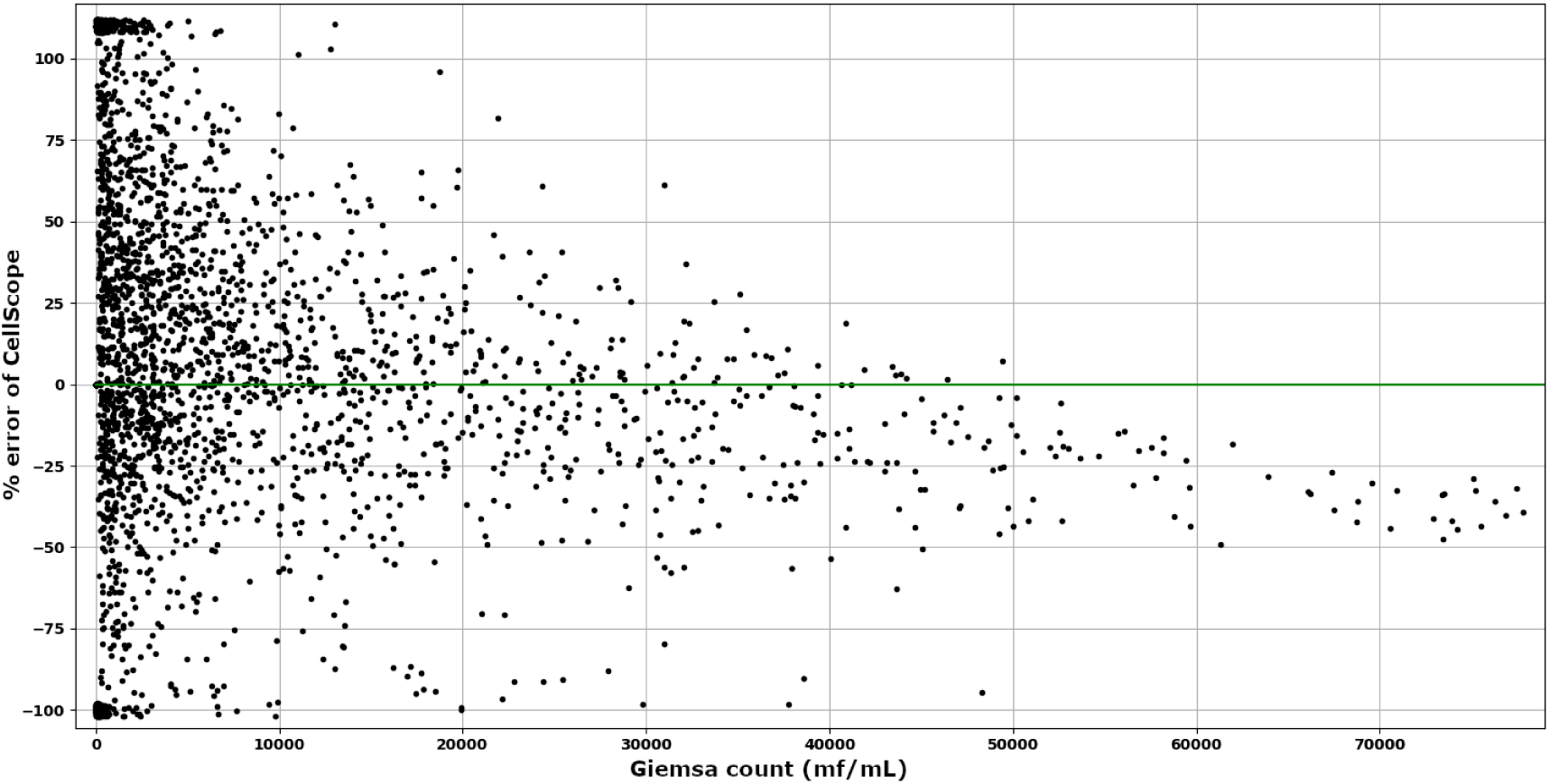
Percent error of historical CellScope vs Giemsa. x-axis = Giemsa count. y-axis = percent discrepancy of CellScope count relative to Giemsa. The extreme undercounts are likely due to cases with undetected coagulation. This is the same data as in **Fig 1E**, visualized differently. Dot locations are jittered, and errors > 100% (including positive counts on negative samples) are bundled at 110%. **Alt Text:** Scatterplot showing CellScope error vs Giemsa ground-truth count. The error is up to +/-70% for Giemsa counts above 10,000 mf/mL.

## Notes

https://zenodo.org/records/21517727

